# A correlation between Long noncoding RNA and unpaired DNA silencing in Drosophila

**DOI:** 10.1101/063677

**Authors:** Manika Pal-Bhadra, Indira Bag, Sreerangam N.C. L.V Pushpavalli, Avadhesha Surolia, Utpal Bhadra

**Author notes:** **Corresponding authors:** Utpal Bhadra, Functional Genomics and Gene Silencing Group, Centre for Cellular and Molecular Biology, Uppal Road, Hyderabad-500007, India. Manika Pal Bhadra, Centre for Chemical Biology, Indian Institute of Chemical Technology, Uppal Road, Hyderabad 500 007, India.

## Abstract

Hybrid transgenes are often recognized as foreign genetic material by cell surveillance mechanisms and are repressed in expression inversely to their copy numbers. Here, we compare the expression of multiple *Adh*-promoter-*white* reporter (*Adh-w*) inserts in paired and unpaired configurations in *Drosophila* somatic cells. The unpaired copies exhibit a clear repression at the transcriptional level relative to paired gene dosage effect, which is dependent upon long noncoding RNA, *Polycomb* and *piwi*. Deficiency mapping using *Adh-w* constructs showed that a minimal sequence of 532 bp of the *Adh* promoter is required for unpaired DNA silencing. Long noncoding RNA detected from this region of the *Adh* promoter is abundant in the unpaired condition. It serves as a docking site for at least two proteins POLYCOMB and Piwi that are essential for active transcriptional silencing. The lesser abundance of noncoding RNAs in the paired configuration only allows PC binding. An active RNA-Protein complex binds to unpaired copies. The loss-of-function *piwi* mutation relieves transcriptional silencing even in association with POLYCOMB. It suggests that functional RNA-Piwi complex might create a silencing driven chromatin configuration by accumulating histone modifying enzymes at the *Adh-w* promoter target. This distinct transcriptional silencing that is stronger for unpaired DNA represents a novel mechanism to repress new transposon and foreign DNA insertions for protection of genome integrity.

## INTRODUCTION

Silencing mechanisms in eukaryotic cells operate at both the posttranscriptional and transcriptional level as stable control mechanisms against transposable elements, virus's foreign hybrid DNA etc. They have also been co-opted to establish functional chromatin domains within chromosomes such as heterochromatin (Volpe et al., 2002, Moazed, 2009, Pal-Bhadra et al., 2002b, Pal-Bhadra et al., 2004a, Haynes et al., 2007, Fagegaltier et al., 2009, Berry et al., 2009). These processes are heterogeneous in mechanism but have common small regulatory RNAs that mediate the silencing (Grewal and Elgin, 2007). Some mechanisms act in cis at the site of RNA production while others operate in trans. These include posttranscriptional and transcriptional silencing both (Aravin and Tuschl, 2005).

Silencing of transposons is mediated by an endogenous RNAi pathway to produce small interfering RNAs (siRNAs) that recognize homologous mRNAs and target them for destruction (Chung et al., 2008). In other cases, clusters of retrotransposons produce transcripts that trigger destruction of all homologous RNAs using piRNAs that are bound to Argonaute proteins of Piwi family (Brennecke et al., 2007, Gunawardane et al., 2007, Malone et al., 2009, Aravin et al., 2007, Ghildiyal et al., 2008). The Piwi-piRNA pathway has been commonly perceived as germline specific. Recent studies have begun to explore this pathway in somatic cells (Ross et al., 2014). These studies illuminate multifaceted somatic functions not only in transposon silencing, but also genome rearrangement and epigenetic programming. Piwi expression and piRNA biogenesis in specific somatic tissues demonstrate their diverse functions.

Drosophila Piwi is expressed during early embryogenesis suggesting a critical function of *piwi* in the early stage of somatic development (Ross et al., 2014). Long piRNA precursors are transcribed from specific genomic loci known as piRNA clusters, cleaved and modified in the cytoplasm, and then transported to the nucleus in complex with Piwi. The *flamenco* locus encodes a long single stranded piRNA precursor that is antisense to *flamenco*'s component transposons.

The accumulation of a set of chromatin proteins including POLYCOMB at the silenced sites mediates transcriptional modulation (Pal-Bhadra et al., 1997). Piwi is a nuclear protein of Argonaute family (Cox et al., 2000) that is distinct from the *Polycomb* Group (PcG) complex. Noncoding RNAs from regulatory sequences of *piwi* were found to be involved in nuclear clustering of PcG targets for silencing in trans and are involved in Pc recruitment at the silenced transgene sites (Grimaud et al., 2006) suggesting that Piwi and associated long non coding RNAs might guide PcG to the silenced sites.

Here we describe the phenomenon of somatic silencing by unpaired DNA in *Drosophila* that operates at the transcriptional level. The silencing is directly coincident with noncoding RNA-Piwi protein complex that leads to repressive chromatin architecture on the silenced transgenes studied. The silencing is relieved by mutations in *Polycomb* and *piwi*, which have been previously implicated in various silencing processes (Pal-Bhadra et al., 1997, Pal-Bhadra et al., 2002a, Pal-Bhadra et al., 2004b). Silencing by unpaired DNA has been documented in meiosis in *Neurospora* (Shiu et al., 2001), *C. elegans* (She et al., 2009) and humans (Baarends et al., 2005). It has been studied most thoroughly in *Neurospora* where the mechanism is posttranscriptional (Hynes and Todd, 2003). Genes or transgenes without a pairing partner during prophase of meiosis will trigger the degradation of all homologous copies in the genome that are either paired or unpaired (Shiu and Metzenberg, 2002).

*Drosophila* somatic cells exhibit homologous pairing of chromosomes and thus are susceptible to a similar type of mechanism. However, in this case, the silencing is transcriptional and chromatin based. Such silencing would serve as a novel mechanism to silence transposons with newly transposed copies that would necessarily be present in an unpaired state. Recognition by the cell of this condition triggers a trans-acting transcriptional silencing in somatic cells to reduce the expression of the homologous copies of a transposon family as a means of cellular defense against additional transpositions.

In this study, hybrid transgenes involving the regulatory regions of the *Alcohol dehydrogenase* gene and the structural part of the *white* eye color gene exhibit strong silencing in genotypes with multiple and unpaired copies, showing a dramatic departure from a linear response to copy number under a paired condition. The difference in silencing between paired and unpaired copy number of trangenes provides an ideal platform for assaying the genome defense mechanism against transposon or foreign genetic intruders. This silencing requires functional gene products of the POLYCOMB repressive complex and the Piwi Argonaute family member (Pal-Bhadra et al., 1997, Pal-Bhadra et al., 2002a). Both paired and unpaired multiple copies are associated with *Polycomb*. However, the unpaired state triggers strong association with Piwi-PC complex that leads to EZ histone methyltransferase accumulation resulting in a strong increase of the histone modification H3K27me3. These findings provide a new dimension to gene silencing mechanisms whereby transcriptional repression is fostered by unpaired genes, that silenced all homologous copies in the genome whether paired or not.

## RESULTS

Previously, it was shown that genetic interaction between two non-homologous transgenes *Adh-w* and *w-Adh* was more sensitive to silencing than mutiple *w-Adh* transgenes (Pal-Bhadra et al., 1997, Pal-Bhadra et al., 1999). Two copies of the *w-Adh* transgene at different chromosomal locations reduced the expression of the non-homologous *Adh-w* construct to a level close to the *white* null (Pal-Bhadra et al., 1999). It may be assumed that *Adh* promoter-w reporter genotype as a foreign intruder. The *Adh-w* construct contains 1.8 kb of the *Adh* regulatory sequence upstream from the transcriptional start sites and 6.5 kb of the *white* structural region required for the normal expression of the *white* mRNA (Figure 1A).One copy of the *Adh-w* transgene is expressed several fold higher relative to the endogenous *white* mRNA in a wild type background (Pal-Bhadra et al., 1999) because the *Adh* promoter is stronger. However adult eye color is *Adh-w* is reduced consistently than endogenous *white* as described earlier (Pal-Bhadra et al., 1999). In this study, we mobilized the *Adh-w* construct from cytological position 16B to new genomic locations using the *P* element stock *Δ2–3* (Robertson et al., 1988). The site of insertion was already identified by *in situ* hybridization (Pal-Bhadra et al., 1999). Three *Adh-w* single insertion lines that exhibit a minimal level of positional variation were used for further study. The neighboring sequence of each *Adh-w* insert was determined by employing inverse PCR (Sullivan et al., 2000). The cytological locations of the two autosomal inserts were 55C on chromosome 2 and 70C on chromosome 3. Using multiple genetic crosses, these single inserts were combined to generate a series of stocks carrying one to six *Adh-w* copies with varying numbers at these three independent locations. The X chromosome of all stocks was homozygous for an endogenous *white* null allele, *w^67c23^*; thus, *Adh-w* transgenes are the sole source of the *white* mRNA and eye color pigments. Single *Adh-w* inserts in the *white* minus background are expressed normally but differ slightly due to minimal positional variation. The synchronized adult transgenic flies of the same age were used for eye pigment assay. The amount of the eye pigment showed a gene dosage effect when each *Adh-w* insert was made homozygous to form a paired configuration. Interestingly, two *Adh-w* transgenes that are heterozygous in two separate locations showed reduced expression in the adult eyes (Figure 1B, C; **S1**). The amount of eye color pigment of two copies of *Adh-w* in a hemizygous condition, as presented in the bar diagram, was less than the single *Adh-w* transgene inserted at either location (Figure 1C). These results imply that expression of two *Adh-w* inserts at different locations is susceptible to more silencing, while paired copies shows a dosage effect.

**Figure 1.**
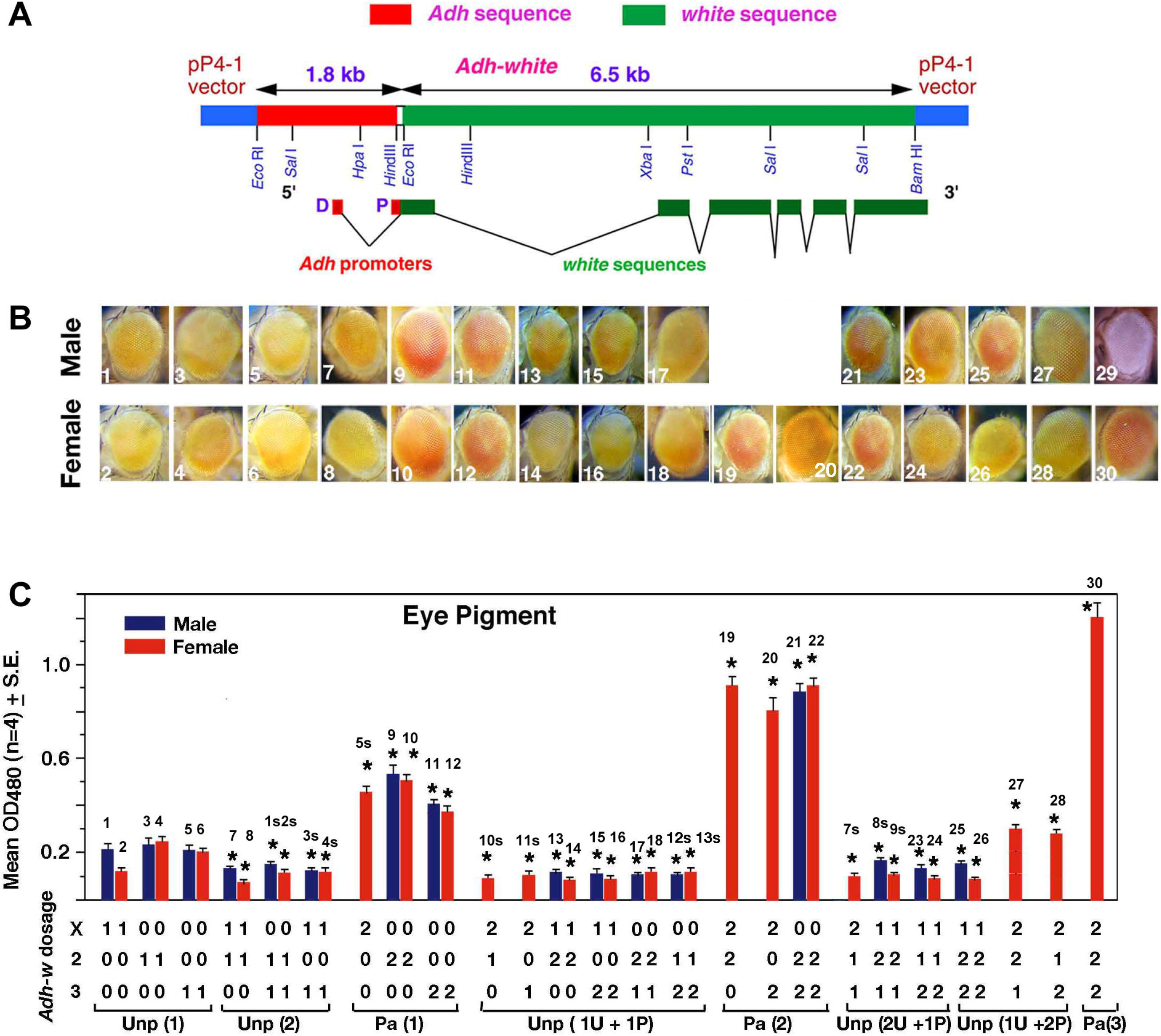
Effect of *Adh-w* copy number on silencing. (A). Structure and restriction map of the *Alcohol dehydrogenase* promoter-*white* reporter construct. The exon and intron boundaries of the *white* coding region and the direction of transcriptional orientation are displayed by arrows (modified from Pal-Bhadra et al., 1999). (B). The eye color of *Adh-white* flies carrying an unpaired copy shows greater reduction relative to paired copies. All flies are in a *y w ^67c23^* background. The genotypes are as follows 1) *Adh-w#1/Y*; +/+;+/+ 2) *Adh-w#1*/+; +/+; +/+ 3) *+/Y; Adh-w#2*/+; +/+ 4) *+/+; Adh-w#2/+; +/+* 5) *+/Y; +/+; Adh-w#3/+* 6) *+/+; +/+; Adh-w#3/+* 7) *Adh-w#1/Y; Adh-w#2/+; +/+* 8) *Adh-w#1/+; Adh-w#2/+, +/+* 9) *+/Y; Adh-w#2/Adh-w #2; +/+* 10) *+/+; Adh-w#2/Adh-w #2; +/+* 11)*+/Y; +/+; Adh-w#3/Adh-w#3* 12) *+/+; +/+; Adh-w#3/Adh-w#3*13) *Adh-w#1/Y; Adh-w #2/Adh-w #2; +/+* 14)*Adh-w#1/+; Adh-w#2/Adh-w#2; +/+* 15) *Adh-w#1/Y; +/+; Adh-w#3/Adh-w#3* 16) *Adh-w#1/+; +/+; Adh-w#3/Adh-w#3* 17)+/+; *Adh-w#2/Adh-w #2; Adh-w#3/+*18) *+/+; Adh-w#2/Adh-w#2; Adh-w#3/+*19) *Adh-w#1/Adh-w#1; Adh-w#2/Adh-w#2; +/+* 20)*Adh-w#1/Adh-w#1; +/+; Adh-w#3/Adh-w#3*21) *+Y; Adh-w#2/Adh-w#2; Adh-w#3/Adh-w#3* 22)+/+; *Adh-w#2/Adh-w#2; Adh-w#3/Adh-w#3* 23) *Adh-w#1/Y; Adh-w#2/Adh-w#2; Adh-w#3/+* 24) *Adh-w#1/+; Adh-w#2/+; Adh-w#3/Adh-w#3* 25) *Adh-w#1/Y; Adh-w#2/+; Adh-w#3/Adh-w#3* 26) *Adh-w#1/+; Adh-w#2/Adh-w#2; Adh-w#3/Adh-w#3* 27) *Adh-w#1/Adh-w#1; Adh-w#2/Adh-w#2; Adh-w#3/+*28) *Adh-w#1/Adh-w#1; Adh-w#2/+; Adh-w#3/Adh-w#3*29) *yw^67c23^/Y* 30) *Adh-w#1/Adh-w#1; Adh-w#2/Adh-w#2; Adh-w#3/Adh-w#3*. (C) The abundance of eye pigments (OD 480) in male and female adult eyes is depicted in the bar diagram. Values calculated for bar diagram were depicted from eye colour of adult flies which were showed in Figure 1B and Figure S1. Values are the mean of three independent pigment assays. 48 heads were used for each assay of each genotype and sex. The number of each genotype corresponds to those in B. Error bars delimit the 95% confidence interval. The values noted by asterisks are statistically different from one *Adh-w* copy male and female values. Thenumber of adults eyes (1–30 in Figure 1B. and (1s–12s in Figure S1) were noted at the top of each bar.

To establish further the effect of transgene copies in paired and unpaired configurations, a series of *Adh-w* stocks were generated that contained various combinations of *Adh-w* copies in paired and unpaired conditions in three different locations eventually producing a stock with a maximum of six copies that are paired in the three separate locations. In different sets of females carrying three *Adh-w* copies either hemizygous at three independent locations or in a combination of a paired set and an unpaired copy elsewhere, the amount of pigment was reduced to an extent close to the two unpaired copies (Figure 1B, C). Thus, in three copies fly, one unpaired copy was able to reduce the expression of one paired copies in trans. When one more copy was added to produce four *Adh-w* copies in a combination of two hemizygotes at different sites and a paired set at one location versus four copies from two paired sets, the former showed less expression than the latter two paired copies. In comparing paired sets, the level of eye pigment increased with the copy number although not strictly proportionally (Figure 1C). In contrast, two five copy *Adh-w* stocks in which one unpaired copy is present together with two paired copies, an amount of pigment higher marginally than two unpaired copies was found. Addition of a single transgene to five *Adh-w* copies that converts them to an all paired configuration resulted in a dramatic increase of the adult eye pigment. The six copies that are paired in all three insertion sites showed a negligible level of reduction relative to the predicted gene dosage effect (Figure 1B, C). Thus, the presence of only one unpaired copy is sufficient to repress two paired copies. The maximum level of repression, as found in this material, is distinct from the *w-Adh* transgene cosuppression (Pal-Bhadra et al., 1997) and *Adh-w/w-Adh* non-homologous cosuppression (Pal-Bhadra et al., 1999), in which the amount of silencing is inversely proportional to the copy number. In this case, the magnitude of silencing is greatly influenced by the paired or unpaired configuration of the hybrid genes.

### Unpaired DNA promotes *Adh-w* transcriptional silencing

To test whether the effect of unpaired *Adh-w* transgenes on silencing is transcriptional or posttranscriptional, we examined the steady state level of the *Adh-w* transcripts using northern blot hybridization and estimated the rate of transcription by measuring the nascent RNA through transcriptional run on assays from the same genotypes. The level of *white* mRNA from two unpaired *Adh-w* copies was reduced, while from the paired copies, *white* transcripts increased proportionately to their copy number (Figure 2, **Top panel**) in two separate locations. A significant reduction of the *white* mRNA in the presence of unpaired copies in different locations and normal expression of the same transcripts in paired configuration were closely matched to their eye pigment assay (Figure 2, **Top panel**). To ensure homogeneity, the relative values in one *Adh-wcopy* in all three locations were close to two folds produced by the paired copies (**Figure S1**).

**Figure 2.**
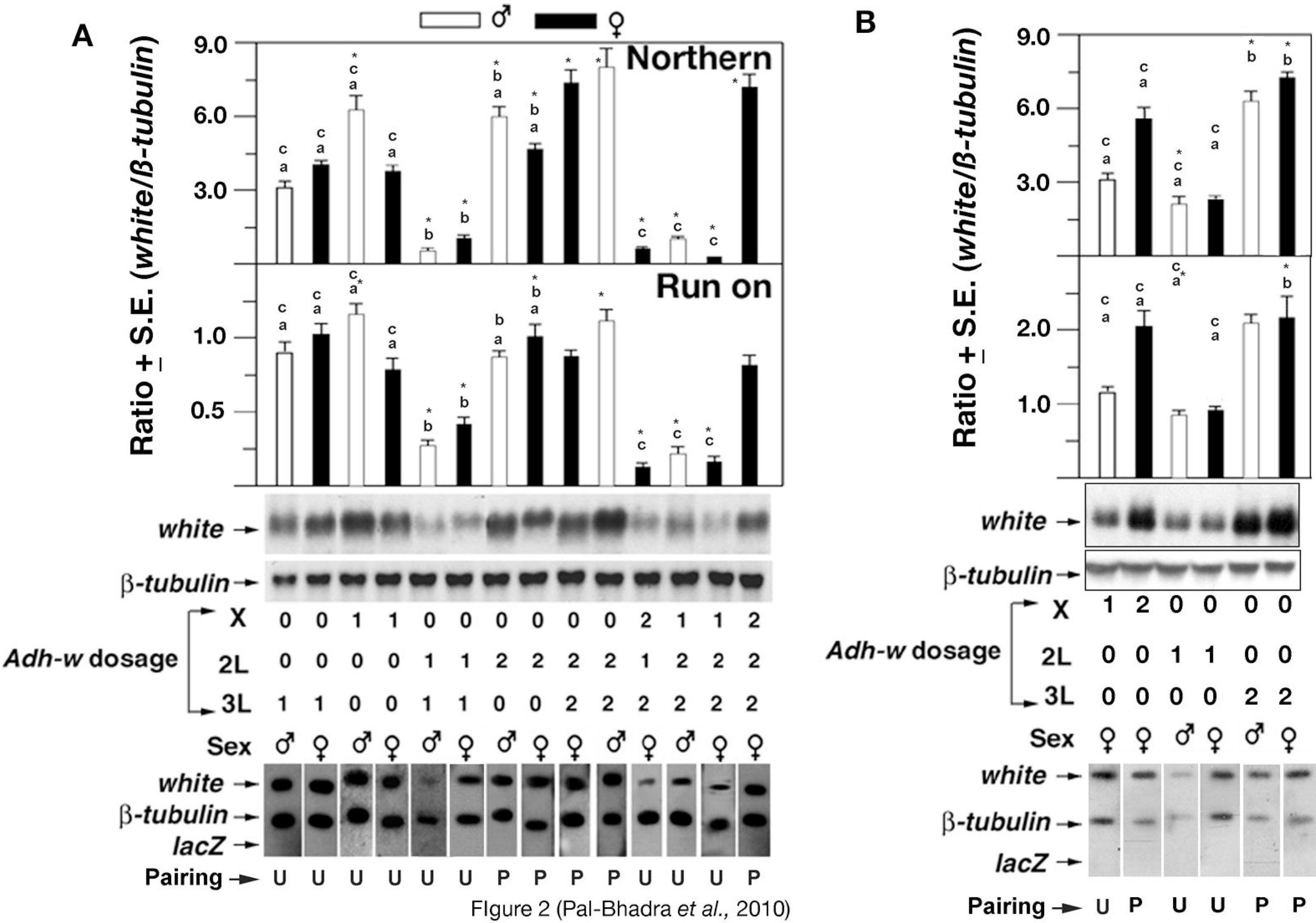
Paired and unpaired *Adh-w* silencing is transcriptional. Autoradiograms of Northern blot hybridizations and nuclear run-on of different classes of adult flies carrying zero to six doses of *Adh-w* hybrid genes probed with *P^32^* radiolabelled antisense *white* RNA are shown. Measurement of *white* transcripts was performed from triplicate experiments. A similar graph was plotted using *white/β-tubulin* ratios of northern gels using the same genotypes. Black bars represent females and open bar represents males. The chromosomal locations, copy number of the *Adh-w* copies and sex of each genotype are noted in the middle of the panels. Mean values marked with an asterisk from triplicate determinations are noted in comparison with the values for the respective one copy *Adh-w*/+ control males and females, respectively, that are significantly different at the 95% confidence level. Pairing configuration of *Adh-w* transgenesis is noted as U-unpaired and P-paired. Values marked with “a” are compared between one copy of *Adh-w* and two paired copies. Values marked with “b” are compared between two paired and two unpaired copies. The bars marked with “c” is compares a single *Adh-w* copy with one unpaired and two paired copies.

By performing nuclear run on assays, the rate of *Adh-w* transcription was measured by estimating the amount of *white* mRNA produced by these transgenes. In these geneotypes the *Adh-w* is the only source of *white* mRNA as the stocks are produced in a *white* null background that does not produce any functional transcripts due to deletion. Here, *β-tubulin* transcripts serve as an internal control while *lacZ* served as negative control. In each genotype, the rate of transcription was also reduced in the presence of an unpaired transgene, while paired copies exhibited higher transcriptional rates. The transcriptional rate of two unpaired (hemizygous) copies was reduced markedly relative to the rate of a single paired copy. In extreme cases (5 copy flies), one unpaired copy is sufficient to reduce the accumulation of *white* transcript produced by two paired copies in different genomic locations (Figure 2, **Bottom**).

Comparing the relative ratios between *white* nascent RNA and *β-tubulin* RNA measured by transcriptional run on and steady state level of RNA estimated by northern hybridization showed that presence of a single unpaired copy promotes silencing, while paired copies are only silenced in a trace level (Figure 2). A parallel reduction of nascent and mature RNA revealed that copy number in *Adh-w* transgenes involves a novel type of transcriptional silencing in which unpaired DNA triggers strong silencing in somatic cells.

### A dosage effect of full-length *white* transgenes

To examine whether *Adh-w* silencing by the unpaired transgene DNA is mediated by the common upstream sequence of the transgenes or by the shared *white* coding regions, we constructed stocks carrying one to six copies of a full length *white* transgene. Each construct is 10.2 kb in length in which a 3.7 kb region upstream to the transcriptional start site is present with 6.5 kb of the *white* protein coding sequences(Hazelrigg et al., 1984). The promoter fragment contains all the regulatory domains including the eye enhancer that is required for normal *white* expression in the larval eye imaginal discs, malpighian tubules, adult eyes etc. In contrast, as noted above, the *white* coding region in the *Adh-w* transgene is placed downstream of the 1.8 kb *Adh* regulatory region. Although the protein coding region in the two transgenes is the same, their regulatory sequences do not share any homologous domain. A northern blot analysis using total cellular mRNA extracted from normal male and female individuals having one to copies of the full length *white* transgene revealed that the mature mRNA was accumulated in a dosage sensitive manner (Figure 3A). Similarly to *Adh-w* transgene dosage, the expression of *white* RNA was greater in males relative to the same copy number in females. These results indicate that silencing by an unpaired transgene maintains the status quo of sex chromosomal dosage compensation and is restricted to the hybrid *Adh-w* construct. Although *Adh-w* and *white* full length constructs share the entire *white* protein coding sequence, silencing does not occur for the full-length *white* construct. The other parental gene of the *Adh-w* hybrid constructs. Endogenous *Adh* transgenes showed a gene dosage effect when one to five copies of *Adh* were examined earlier (Pal-Bhadra et al., 2002a).

**Figure 3.**
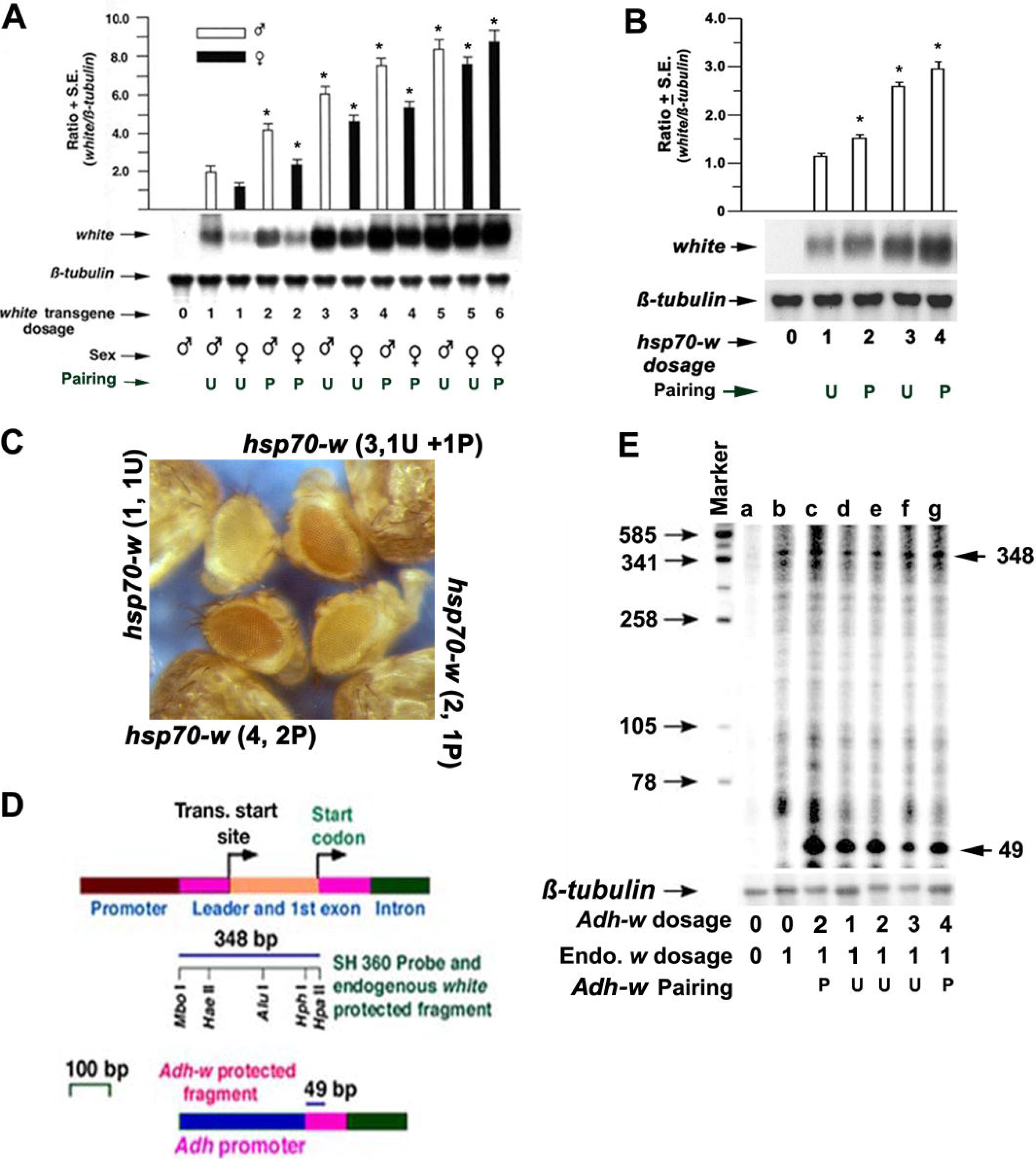
Paired and unpaired *Adh-w* transgene silencing is promoter dependent. **(A)** A dosage dependent expression of zero to six full length *white* transgene copies. Representative Northern autoradiogram using adult RNA isolated from flies carrying one to six copies of full-length *w* transgenes with no endogenous *white* gene is shown. The means of three independent *white/β-tubulin* ratios are represented by the bar diagrams. The values marked by an asterisk are significantly different from the respective controls at the 95% confidence level. The number of *w* transgene copies and sex of the flies are noted at the bottom. U-unpaired, P-paired. **(B)** Autoradiogram showing the northern hybridization of the *white* transcript levels was dependent on *hsp70-w* transgene dosage. The *white/β-tubulin* ratios marked with asterisks are significantly different from the one copy *hsp70-white* control level. The number of *hsp70-w* copies is noted at the bottom. U-unpaired and P-paired. **(C)** A similar gene dosage effect on eye pigment was found in adult female flies carrying zero to four *hsp70-w* transgenes. The intensity of pigment accumulated in the adult eye was proportional to the *hsp70-w* dosage. The *hsp70-w* copy number is noted in the parentheses. **(D)** Expected products from RNAse mapping experiments using the *white* SH360 probe. The primary transcripts of the *white* gene are illustrated at the top. The promoter and leader sequence of the first exon and intron are noted. The length and restriction map of the SH360 probe are indicated by a line below. The RNA splicing pattern and protected fragments of endogenous *white* and *Adh-w* transcripts are depicted. The *Adh-w* transcripts also give rise to a 348 bp protected fragment. The total abundance of endogenous *white* transcripts is represented by the 348 bp fragment. **(E)** RNAse protection assay of *white* mRNA. The RNA from different classes of adult females carrying zero to four *Adh-w* transgenes was estimated. Lanes are: (a) *w^1118^/w^1118^*, (b) *+/w^1118^*, (c) *Adh-w#1/w^1118^*, (d) *Adh-w#1/+*, (e) *Adh-w#1/+; Adh#3/+*, (f) *+/w^1118^; Adh-w#2/+; Adh-w#3/Adh-w#3*, (g) *+/w^1118^; Adh-w#2/Adh-w#2; Adh-w#3/Adh-w#3*. Predicted size of the protected fragments based on the RNA splicing pattern is indicated on the right and the sizes of the marker RNAs are shown. A 70 bp protected fragment of *β-tubulin* RNA (separate panel) was hybridized with the same blot as a gel loading control. The pairing configuration of *Adh-w* transgene copies were noted as U-unpaired and P-paired, respectively.

### *Adh-w* silencing is context specific

To define whether the *Adh* promoter alone or hybrid sequences at the promoter-reporter junction is necessary for unpaired DNA promoted cosuppression, *ahsp70-white* hybrid construct was examined. In this transgene, a 900 bp long inducible promoter of the heat shock protein 70 is fused to the *white* coding sequences and is present in a *white* minus background (Steller and Pirrotta, 1985). In this stock *white* mRNA is expressed at a constitutive, minimal level at the normal 25°C temperature but is over-expressed by heat shock at 37°C. Two separate *hsp70-white* inserts located at different autosomal locations (chromosomes 2 and 3) were combined to generate four copy stocks. We measured the eye pigment as well as *white* mRNA of the adult flies carrying one to four copies. The amount of pigment level is proportionally elevated with the increasing number of the *hsp70-w* dosage irrespective of their unpaired or paired configuration (Figure 3B, C). These results were further verified by estimating *white* mRNA level from the same genotypes. The accumulation of *white* transcript showed a gene dosage effect (Figure 3B, C). Earlier findings showed that full-length *Adh* transgenes showed a dosage sensitive effect proportional to the copy number increment within this range (Pal-Bhadra et al., 2002a). These results reveal that another promoter-white reporter junction is not sufficient for initiating unpaired DNA promoted cosuppression. Therefore, the context of the *Adh* promoter-*w* reporter sequence is implicated as necessary for silencing.

### No influence on endogenous *white* expression

To determine whether cosuppression in the *Adh-w* dosage series that is triggered by the unpaired copies has any influence on the endogenous *white* expression, we combined an endogenous *white* copy in its normal location with one to four copies of the *Adh-w* transgenes in two different autosomal locations. In northern blot hybridization analyses, *white* mRNA levels contributed by the endogenous *white* locus cannot be differentiated from the product of the *Adh-w* transgenes in the combination flies, as there is no difference in the size of the *white* mRNA derived from the endogenous *white* and the *Adh-w* transgenes. However, in the *Adh-w* construct, the *Adh* promoter is joined to the *white* coding regions preceding the translational start codon and including the *white* leader sequences (Figure 3D). The truncation of the 5’ end of the native *white* mRNA produced by the *Adh-w* construct provides an opportunity to distinguish them from the transcripts synthesized by the endogenous *white* gene. We used a quantitative RNase protection assay that distinguishes the endogenous *w* mRNA and *Adh-w* mRNAs to estimate their products separately. The protecting fragment recognizes 340 bp of RNA from the endogenous transcripts and a 49 bp fragment of *Adh-w* (Figure 3E). As a gel loading control, a 70 bp fragment complementary to the *β-tubulin EcoR1-Sau3A* fragment was protected by simultaneous inclusion in the same reaction. The results reveal that protected fragments of the transgene were strongly reduced in the presence of an unpaired copy in the genotype. In contrast, the fragment protected by the endogenous transcripts was equally expressed in all genotypes irrespective of the *Adh-w* copy number (Figure 3E, Table 1). These results suggest that multiple copies of the *Adh-w* transgene have no influence on the endogenous *w* mRNA in trans implying that a common coding sequence between endogenous *white* and the *Adh-w* transgenes is not sufficient for silencing to occur.

**Table I.** Measurment of band densities of endogenous *white* band and *Adh-w* band from triplicate RNA protection assay gels relative to *β-tubulin* bands.

| ***Adh-w***<br>**dosage** | **Endogenous**<br>***w* dosage** | **Paired/unpaired** | **Band intensity**<br>**(348/β-**<br>**tub)+S.D.** | **Band**<br>**intensity**<br>**(49/$\beta$ tub)**<br>**+S.D** |
| --- | --- | --- | --- | --- |
| 0 | 0 |  | 0.07 + 0.003 | - |
| 0 | 1 |  | 1.53 + 0.04 | 0.05 + 0.001 |
| 2 | 1 | Paired | 2.83 + 0.09 | 9.39 + 0.7 |
| 1 | 1 | Unpaired | 1.49 + 0.04 | 7.51 + 0.6 |
| 2 | 1 | Paired | 1.31 + 0.03 | 7.76 + 0.5 |
| 3 | 1 | Unpaired+<br>Paired | 1.79 + 0.05 | 3.04 + 0.2 |
| 4 | 1 | Two Paired | 1.91 + 0.03 | 7.87 + 0.5 |

To test the effect further in the adult eye, we compared eye pigment amounts in adult *w/+* female flies carrying either zero or four copies of the *Adh-w* transgenes. The presence of the *Adh-w* transgenes does not alter the eye color of the flies, which is consistent with a lack of silencing found for the mRNAs (**Figure S3**).

### Unpaired silencing requires a 532 bp *Adh* promoter sequence

To test whether *Adh* promoter sequence or the junction of the *Adh*-promoter-*white* reporter construct is required for paired and unpaired silencing, we replaced the *Adh-w* construct with another transformant containing the same 1.8 Kb *Adh*-promoter fused with the *Gal4* coding region (Palm et al., 2012) located on chromosome 3. Analysis of the adult flies carrying paired copies of *Adh-w* transgenes showed that addition of one copy of *Adh-Gal4* on chromosome 3 produces a silenced condition (Figure 4). These results suggest that the 1.8 kb *Adh* promoter fused with different reporter constructs is sufficient to mediate unpaired silencing.

**Figure 4.**
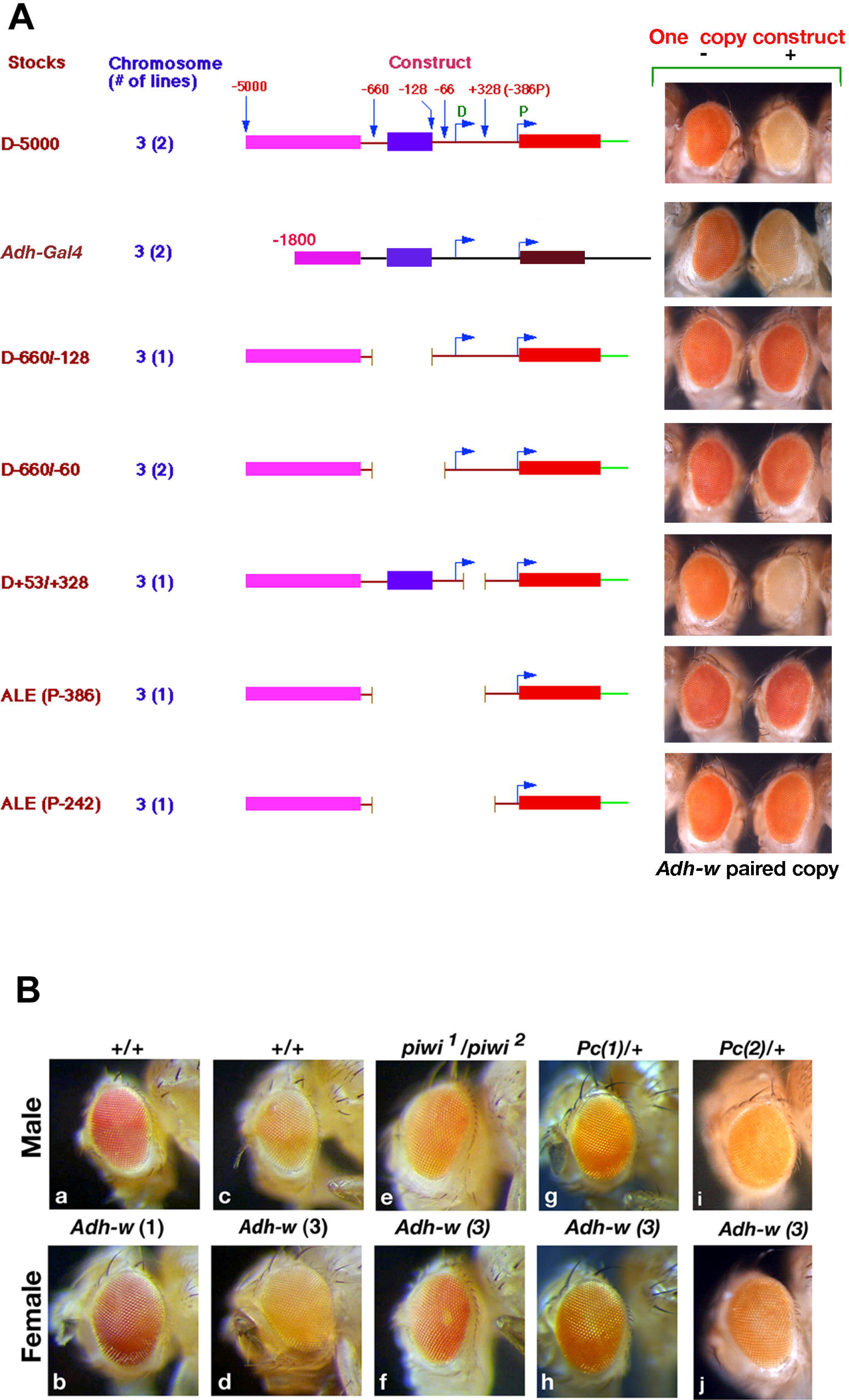
Genetic map for *Adh* promoter sequence required for unpaired DNA silencing. **(A)** The map of a series of *Adh* promoter truncated constructs including full length*Adh-w transgenewas* shown. One copy of each construct and full length *Adh-w* hybrid gene was combined with *Adh-w* paired copies. The interaction without or with the construct in the adult eye was shown in right side of the panel. The stock name and chromosome location are noted at the left. The structure and the deletion of each construct is noted in cartoon structure. **(B)** Eye color of *Adh-w/Y* and *Adh-w/Y, Adh-w/ Adh-w (Adh-w-3)* flies carrying one and three copies of transgenes in a background of normal, heterozygous *Polycomb* or heteroallelic *piwi* mutations. All flies have a *w^-^* background. The mutation and *Adh-w* transgene copy number are noted in the parenthesis.

A series of progressively truncated *Adh* constructs (Corbin and Maniatis, 1989b, Corbin and Maniatis, 1989a) were tested to map the minimal *Adh* promoter sequence needed for unpaired silencing (Figure 4). One copy of each construct was combined with a paired set of *Adh-w* transgenes. The eye color of the *Adh-w* paired copy was examined in the presence of each construct separately. Four of the five constructs failed to elicit unpaired silencing. The mapping of overlapping deletion regions suggested that a 532 bp *Adh* promoter sequence from −660 to −128 bp before the distal *Adh* transcriptional start site is required for unpaired silencing. The same sequence is required for homology recognition of transgene cosuppression (Pal-Bhadra et al., 1999) that includes the adult enhancer region of the *Adh* gene.

### Effect of *piwi* and *Polycomb* mutations on unpaired *Adh-w* silencing

To investigate whether *Adh-w* silencing by unpaired transgene DNA is affected by mutations in *piwi* and *Polycomb* as previously found involved with transgene cosuppression (Pal-Bhadra et al., 1997, Pal-Bhadra et al., 2002a), we produced flies with three copies of *Adh-w* carrying one unpaired transgene with heteroallelic *piwi(piwi^1^/piwi^2^)* and two heterozygyous *Pc* [*Pc(1)/+, Pc(2)/+*] mutations through a series of genetic crosses (See Supplementary Materials). We observed a marked increase in the eye pigment level in the mutational background of *piwi^1^/piwi^2^* and *Pc/+* [*Pc(1)/+, Pc(2)/+*], even in the presence of an unpaired transgene indicating that this silencing was disrupted substantially by the *piwi^1^/piwi^2^* and *Pc/+* [*Pc(1)/+, Pc(2)/+*] genotypes (Figure 4B; **S3**). However, the increase of pigment does not exceed the level produced by a single *Adh-w* transgene copy (Figure 4B). The amount of silencing was further verified at the *white* transcript level using northern blot hybridization. Total cellular RNAs were isolated from different genotypes and hybridized with *white* cDNA probes. The results show that the presence of heterozygous *Pc (Pc/+)* and heteroallelic *piwi (piwf/piwi^2^)* mutations does not alter the *white* mRNA level in *w^67c23^; Adh-w/+* flies relative to the control (**Figure S4**). However, the mutational effect of *piwi* and *Pc* in genotypes with three *Adh-w* copies (one unpaired) shows that the *white* mRNA accumulation increased 61% and 47%, respectively. The effect is stronger in males (59%) compared to females (49%) carrying a copy of the *Pc* [*Pc(1)/+, Pc(2)/+*] mutation relative to control males and females. A similar increment was also found when the *piwi (piwf/piwi^2^)* mutation was used in trans-heterozygotes (**Figure S4**). The presence of one copy of the *Pc* [*Pc(1)/+, Pc(2)/+*] mutation and heteroallelic *piwi* (*piwi^1^/piwi^2^*) mutations reduced the unpaired *Adh-w* silencing markedly.

### Co-localization of Piwi and POLYCOMB proteins on unpaired *Adh-w* transgenes

Piwi is a member of the Argonaute family, which is anticipated to localize mostly to the cytoplasm(Thomson and Lin, 2009). However, earlier studies have shown that Piwi is accumulated in the nuclei in early *Drosophila* embryos and is also visualized in the polytene chromosomes of third instar larvae (Cox et al., 2000, Brower-Toland et al., 2007). We immunostained polytene chromosomes of wild type larvae with a dilution series of commercially available Piwi antibodies (AbCam, UK). The Piwi protein was not detected in most cases when immunostained at standard antibody concentration range (**Figure S5**). However, at higher antibody concentration, Piwi was detected in the chromo-center and many euchromatic sites (Figure 5A). A heteroallelic *piwi (piwf/piwi^2^)* mutant control demonstrated that the level of antibody binding is dramatically reduced but the binding capability on the polytene chromosome was not eliminated (**Figure S5**), which indicates that reduction in chromosomal binding of antibodies is Piwi antibody dependent. Double immunostaining of the polytene chromosomes with anti-PC and anti-Piwi antibodies showed that PC and Piwi bind to numerous euchromatic sites, but PC is more prominent than PIWI. PC labeled sites considerably (61%) overlap with Piwi enriched sites (Figure 5B).

**Figure 5.**
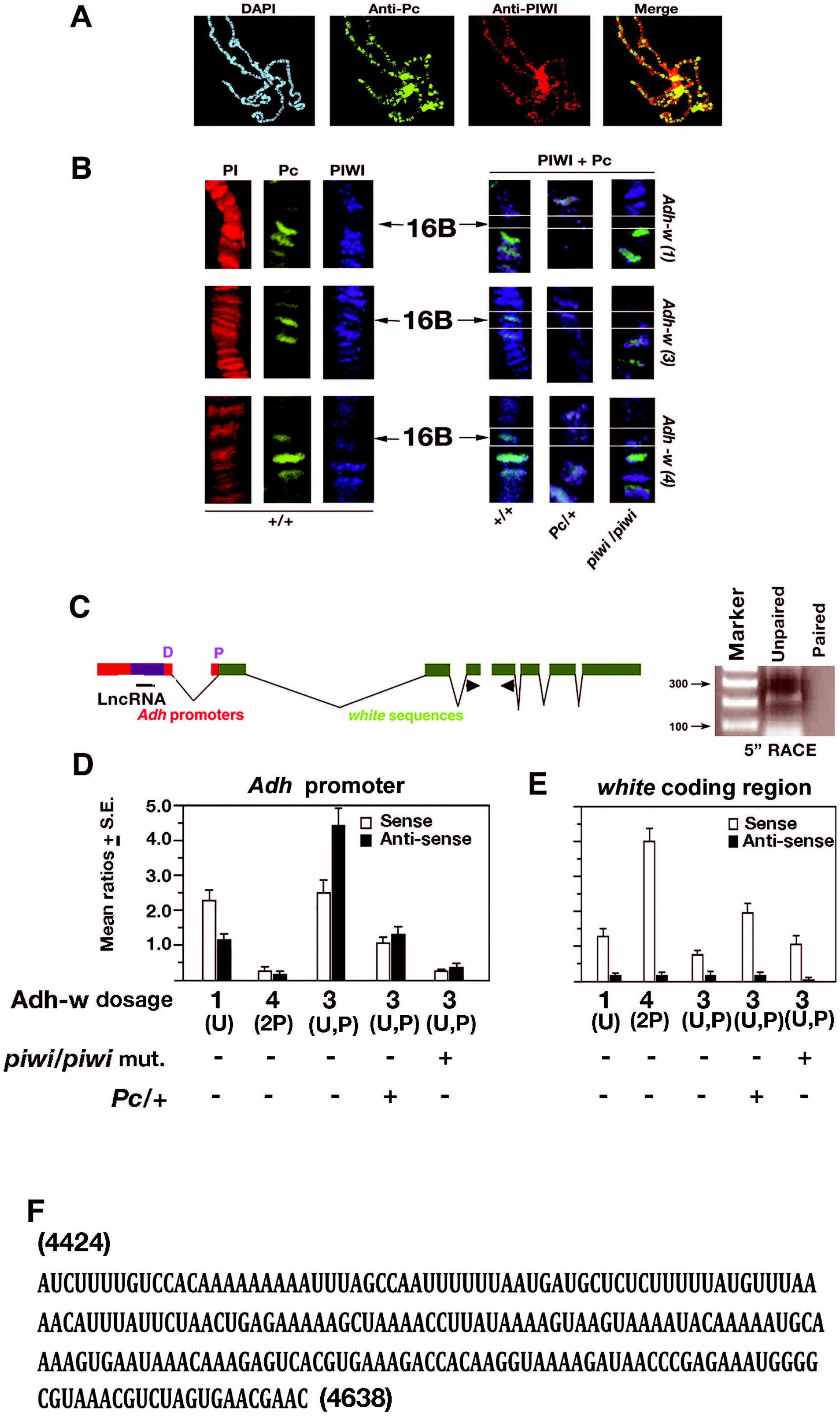
The effect of *piwi* and *Pc* on the *Adh-w* paired and unpaired silencing. **(A)** The overlapping binding of Polycomband PIWI proteins on the giant chromosomes of the third instar female larvae was determined by double immuno-staining with anti-PC (green) and anti-PIWI (red) antibodies. The overlapping image of the PC and PIWI binding is at the right. Merge figure of the polytene chromosome was shown. **(B)** The superimposed images showing localizations of PIWI (green) and PC (light blue) antibodies on the polytene bands at the X chromosome 16B *Adh-w* insertion site when one, three (1 unpaired + 2 paired) or four (2 paired) copies of *Adh-w* are present in wild type (+/+); *Pc*/+ and *piwi* mutant genotypes. Arrows indicate the position of the 16B transgene. The chromosomes were counterstained with propidium iodide (PI). The white boxes on the close up views of the merged chromosomal segments note the insertion site of *Adh-w* transgene at the 16B region and accumulation of Pc (green) and PIWI (dark blue) proteins. **(C)** Schematic diagram of the *Drosophila Adh-w tr*ansgene with *Adh* promoter. Dual *Adh* promoters (D, P), exon and intron were noted. The primer for antisense 5’RACE amplificant was noted (small black arrow). The size of the antisense product was noted in the *Adh* promoter (black line). Amplification of the long antisense ncRNA was performed by unpaired (*Adh-w* 3 copies) and paired (*Adh-w* 4 copies) copies. DNA marker gives the size of the product. **(D) (E)** Two pairs of *mini-w* promoter specific primers encompassing within the antisense regions of the *Adh* promoter and *white* structural gene (*white* 2^nd^ exon, black arrow) were used for strand specific RT-PCR to identify ncRNAs from three *Adh-w* stocks. Only a sense product was amplified from this region whereas for the structural part of the gene in both sense and antisense, a product was generated. Bars represent relative *miniwhite* promoter RNA expression levels normalized to an internal control (*18S* rRNA). Error bars represent mean value of three replicates of S.D. (p< 0.05). **(F)** Sequence of the 214 bp long noncoding RNA from the *Adh* regulatory region.

Because the mutational effect of *Pc* and *piwi* genes reverses the *Adh-w* silencing by unpaired copies, we tested whether Piwi and PC proteins colocalize at the unpaired *Adh-w* site (16B) on the larval polytene nuclei under strong silencing conditions. The specific site for Adh-w insertion was previously determined by in situ hybridization using w antisense probe in Adh-w construct (Pal-Bhadra et al., 1999). Earlier studies have shown that PC antibodies bind nearly 100-150 endogenous sites(Franke et al., 1992, Rastelli et al., 1993). PC and Piwi proteins are not detected at the *Adh-w* (16B) insertion site when only a single construct is present. This result indicates that silencing of a single *Adh-w* transgene does not occur and serves as a control. However, two other *Adh-w* insertions overlap with normal Pc binding (Figure 5A). The immunostaining of the polytene chromosomes was also carried out in three and four copy *Adh-w* stocks in wild type, *piwi^1^/piwi^2^* and [*Pc(1)/+, Pc(2)/+*] mutant backgrounds. In three copy *Adh-w* larvae, both proteins overlap at the 16B (*Adh-w*) site in the wild type background, which exhibits strong silencing. In the [*Pc(1)/+, Pc(2)/+*] genotype, PC binding was basal and Piwi binding was detected but to a lesser extent than in wild type. In the *piwi* mutants, there was no binding of either PC or Piwi. Conversely, in the four copy *Adh-w* stock, which has two paired copies of *Adh-w*, only PC was detected consistently at the *Adh-w* 16B insertion site in the normal background. In the [*Pc(1)/+, Pc(2)/+*] genotype, PC was detected at a low level whereas Piwi was below detectable level (Figure 5B). In the case of *piwi* mutants, neither Piwi nor PC was detected. Thus, PC binding is dependent on the presence of a functional copy of the *piwi* gene product.

### A long noncoding RNA-protein complex is required for unpaired silencing

To examine the specific role of *piwi* on unpaired silencing, we focused on noncoding RNA generated from 532 bp of *Adh* promoter in adult *Drosophila*. We conducted antisense 5’ RACE analysis using a consistent primer at the start of the 532 bp upstream of the *Adh* distal promoter. Antisense orientation of the primer was used for amplification of *Adh* noncoding RNA. To ensure antisense production, the product was amplified with two sets of primers generated by 5’ RACE product (Figure 5C). The same primers were used for sense strand amplification. In the control, the coding region of *white* sequence was amplified (Figure 5E). The results showed that an abundance of antisense RNA relative to sense RNA is accumulated with homology between -322 bp and -128 bp sequence of the *Adh* promoter

To ensure the presence of long non coding RNA, ncRNA was amplified in real time PCR analysis. The mean ratio of triplicate amplification is plotted in a bar diagram. The findings showed an abundance of antisense transcript relative to sense RNA in the Adh promoter. Antisense transcript was more prevalent in the 532 to Adh promoter in the transgene stock carrying three copies of *Adh-w* compared to four copy stock. In both the stocks, Promoter sense RNA was produced in lesser amount compared to Adh promoter antisense RNA (Figure 5D,E).Moreover the accumulation of Adh promoter specific sense and antisense RNA was proportionately reduced in the present of PC and piwi mutation (Figure 5D). It revealed that accumulation of promoter based antisense and sense RNA were piwi and Pc mutant specific. In contrast the amplification of white coding region showed abundant amplification of sense RNA (Figure 5E). The sequencing of the 5’RACE product and RNA sequencing showed that an isolated 214 bp fragment is the probable long noncoding RNA from the *Adh* promoter, which is abundant inbetween -322 to-128 bp of Adh promoters of Adh-w unpaired construct (Figure 5F).

To determine further whether the antisense product of the *Adh* regulator RNA is a precursor cluster of small piRNA, we performed a small RNA northern blot analysis extracting the RNA from unpaired and paired *Adh-w* copies (**Figure S6**). Small RNA northern was used for identification of piRNA cluster. An in silico sequence analysis of *Adh* regulatory RNA does not recognize any piRNA cluster in the 532 bp of the *Adh* promoter. Among unique Piwi associated small RNA, only 8.7% matched with known non coding RNA. Further, in a northern analysis, a gradual increase of the RNA loading from unpaired and paired *Adh-w* combination did not show any indication of small RNA. The gel was probed with antisense 532 bp Adh promoter RNA (**see Figure S6**). Therefore a non canonical function of piwi is important for unpaired DNA silencing

To ascertain any potential association of Piwi protein with long noncoding RNA generated from the *Adh* regulatory sequence, RNA immuno-precipitation was conducted in the four and five copies *Adh-w* transgenic stocks (Figure 6A). Initially, two colocalized proteins Piwi and PC were immuno-precipitated with long noncoding RNA. Amplification of noncoding RNA homologous to the *Adh* promoter generated a product that was greater in magnitude (2 fold) in the unpaired five copies *Adh-w* stocks (Figure 6B) than the level of amplification in the paired four copies. We also tested immuno-precipitated chromatin from *piwi* heteroallelic and Pc heterozygous mutations. In the case of PC heterozygous mutation, the amount of noncoding RNA accumulation was reduced to half compared to wild type. However, the loss of one functional copy reduced the RNA to a trace level in Piwi immuno-precipitated chromatin (Figure 6C,D). In addition, the presence of heteroallelic *piwi* mutation shows no amplified non-coding RNA immuno-precipitated from the Piwi and PC proteins (Figure 6C,D). The mutational analysis revealed that noncoding RNA showed an interaction with Piwi indicating that Piwi plays a role in mediating the interaction of non coding RNA and its associated proteins.

**Figure 6.**
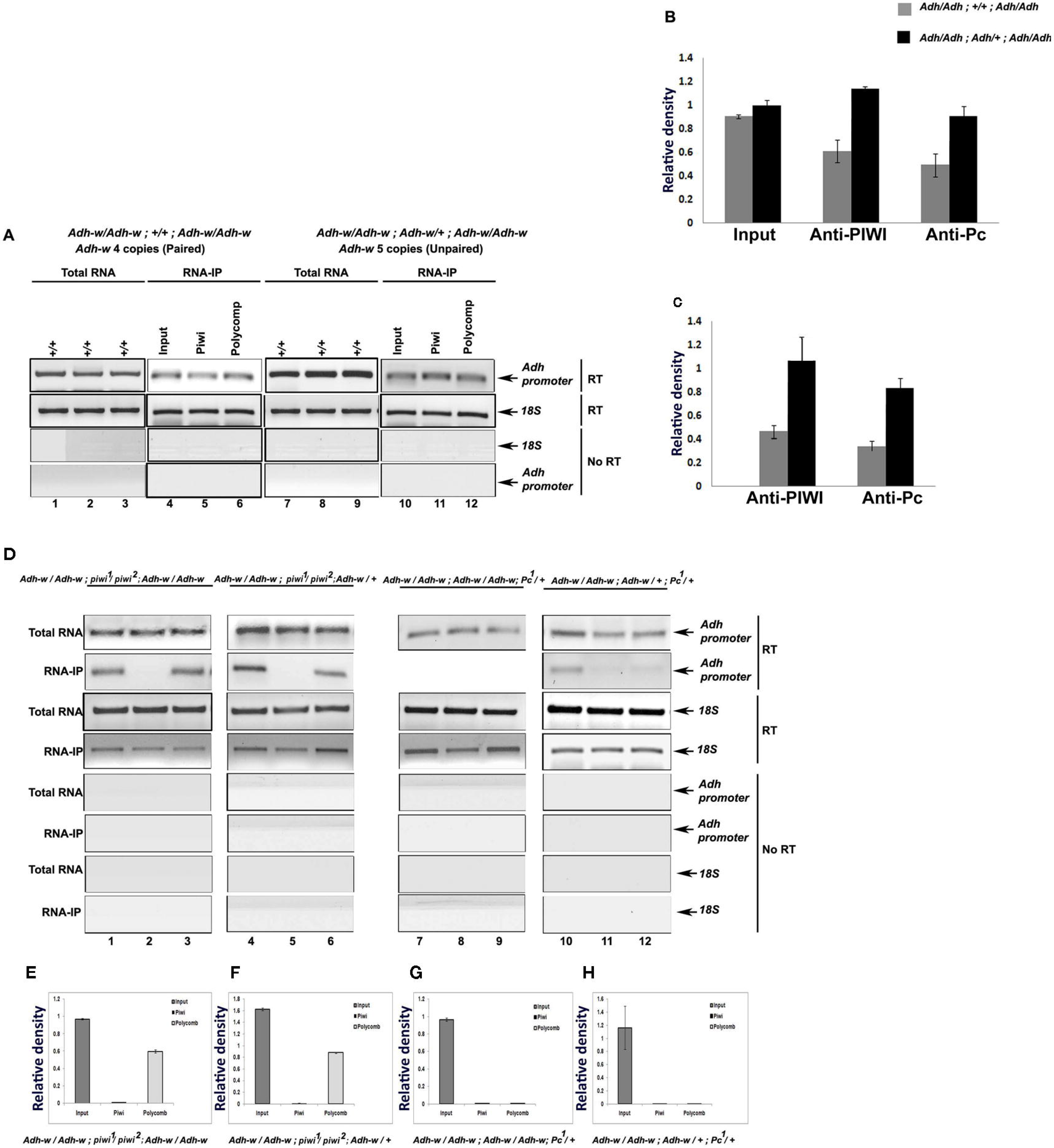
Characterization of long noncoding RNA and RIP assays with anti-PIWI and anti-POLYCOMB antibodies. **(A)** Lanes 1-3 and lanes 7-9 represent total RNA from Drosophila wild type. Lanes 4, 5, 6 represent input, PIWI associated RIP and POLYCOMB associated RIP samples for *Adh-w* 4 copies (*Adh/Adh; +/+; Adh/Adh*). Lanes 10, 11, 12 represent the same as for *Adh-w* 5 copies (*Adh/Adh; Adh/+; Adh/Adh*). **(B)** Bar diagram of relative density of RNA-Immunoprecipitation (RIP) assays was plotted by taking values from three independent experiments performed. *18S* amplification was taken as the internal control. **(C)** Lane 1, 2, 3 represent input sample, PIWI associated RIP sample and POLYCOMB associated RIP sample in *Adh-w* 4 copies with *piwi^1^/piwi^2^* heteroallelic mutation (*Adh/Adh; piwi^1^/piwi^2^; Adh/Adh)*. Corresponding bar diagram of relative density was plotted in **(D)**.**(D)** Lanes 4, 5, 6 represent the same as in **(A)** for *Adh-w* three copies with *piwi^1^/piwi^2^* heteroallelic mutation (*Adh/Adh; piwi^1^/piwi^2^; Adh/+)*. Corresponding bar diagram of relative density was plotted in **E**. **(E)** Lanes 7, 8, 9 represent input sample, PIWI associated RIP sample and POLYCOMB associated RIP sample in *Adh-w* 4 copies with *Pc* mutation (*Adh/Adh; Adh/Adh; Pc^1^/+)*. Corresponding bar diagram of relative density was plotted in **(F)**. **(F)** Lanes 10, 11, 12 represent the same as in (A) for *Adh-w* 3 copies with *Pc* mutation (*Adh/Adh; Adh/+; Pc^1^/+)*. Corresponding bar diagram of relative density was plotted in **(G)**. **(G)** Plot of relative density.

### Association of chromatin factors at the *Adh-w* promoter sequences

A critical question is whether noncoding RNA, Piwi and PC proteins form separate interaction modules with the *Adh* promoter of unpaired *Adh-w* copies with the transcriptional machinery. We assayed the role of noncoding RNA-Piwi interaction in transcription. Because silencing by an unpaired copy of *Adh-w* in the three copies *Adh-w* flies is associated with PC and Piwi recruitment to the insertion sites, the organization of the chromatin on the transgene was analyzed by chromatin immuno-precipitation. The level of repressive Polycomb complex chromatin proteins PC, EZ histone methyltransferase and the histone modification H3K27me3 were estimated at the *Adh* promoter (Figure 7) and *white* exon 2 sequences (**Figure S7**). When one copy of the strongly expressed *Adh-w* transgene is present in the genome, we find only a trace amount of PC, EZ and H3K27me3 at the *Adh-w* promoter, while the same fragment was amplified readily from the whole cell extract (WCE) (Figure 7A). In the strongly silenced three copy *Adh-w* transgene stock, PC and EZ proteins showed a strong accumulation at the *Adh* promoter together with an increased H3K27me3 methylation (Figure 7B) but little change on exon 2 (**Figure S6**). This accumulation of a repressive complex might prevent the binding of RNA polymerase that leads to a strong repression of the *Adh-w* transcripts. In two copyAdh-w stocks with a two unpaired configuration, there is enrichment of the silencing components at the *Adh* promoter allowing restoration of *Adh-w* unpaired silencing (Figure 7C). In four copy stocks with the two paired configuration, only PC proteins are accumulated but there is no EZ and H3K27me3 association suggesting that only Pc mediated binding is applicable (Figure 7D, **S6**). These results suggest that unpaired DNA might trigger a repressive mechanism.

**Figure 7.**
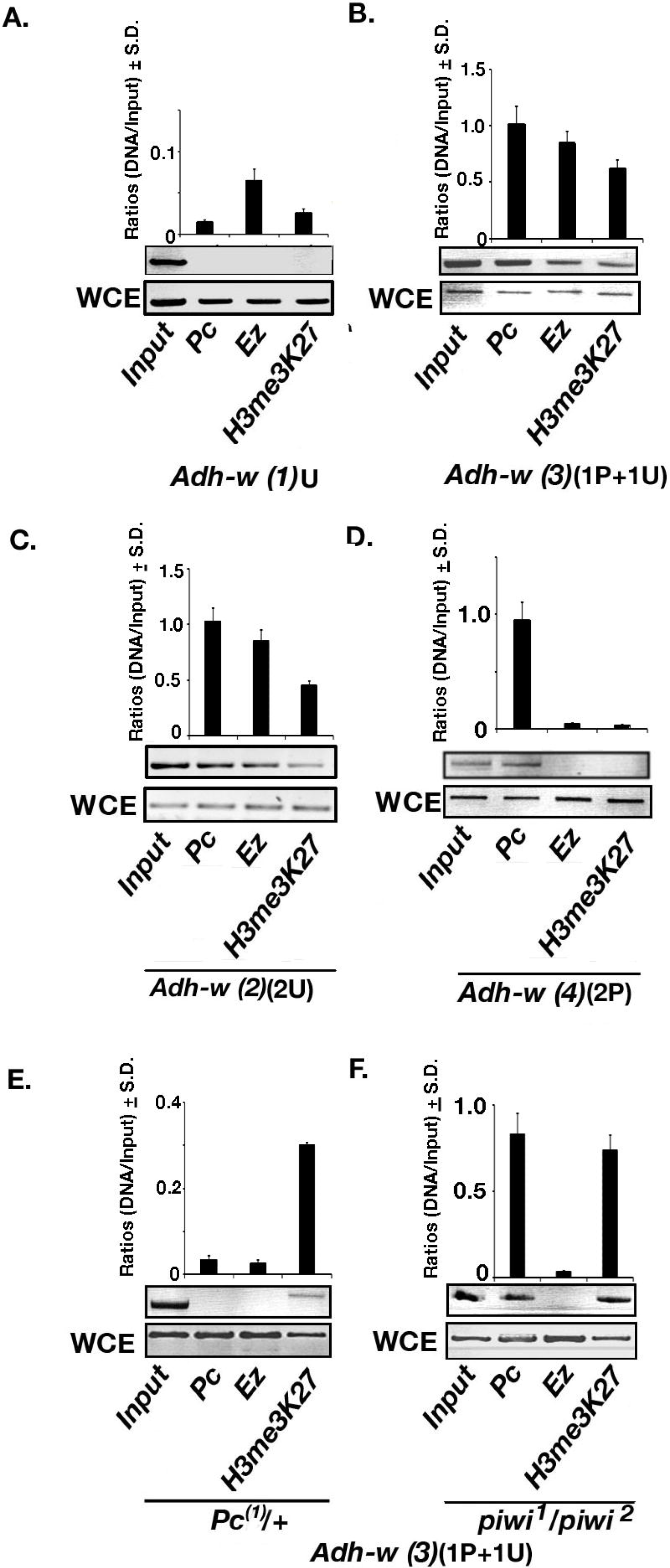
Noncoding RNA-protein complex is required for the transcriptional machinery on the *Adh-w* promoter. Chromatin immunoprecipitation comparing the detection of PC, EZ and H3K27me3 on the promoter of *Adh-w* assayed in third instar larvae of wild type, *piwi* and *Pc/+* mutant strains. The enrichment of proteins for each amplicon of the *Adh-w* locus was measured relative to an amplified fragment by the same primer pairs in whole cell extract (WCE) and the relative ratios from three independent experiments were depicted between each panel. The regions assayed were the distal 532 bp of the *Adh* proximal promoter required for transcriptional cosuppression (Pal-Bhadra et al., 1999). **A)** One copy of *Adh-w* in wild type. **B)** Three copies of *Adh-w* in wild type. **C)** Two (two unpaired) *Adh-w* copies **D)** Four (two paired) copies of *Adh-w*. **E)** Three copies of *Adh-w* in a Pc/+ mutant background. **F)** Three copies of *Adh-w* in a *piwi* mutant background.

ChIP was also performed in genotypes in which [*Pc(1)/+, Pc(2)/+*] or *piwi^1^/piwi^2^* mutations were combined with three *Adh-w* transgenes. As expected, the two Polycomb Group proteins, PC and EZ, are disassociated from the *Adh* promoter sequences in the *Pc/+* mutant with a concomitant drastic reduction of H3K27me3 enrichment (Figure 7E, **S7**). Interestingly, binding of PC and enrichment of H3K27me3 was found in the three copy genotype in the presence of *piwi^1^/piwi^2^* heteroalleles but there was a dissociation of EZ (Figure 7F, **S8**).

## DISCUSSION

We propose the phenomenon of silencing by unpaired DNA in somatic cells of *Drosophila* and provide insight into its mechanism of action. The *Adh-w* transgenes analyzed in multiple copies show dramatic silencing when a single unpaired copy is present in the genome. The silencing operates on the transcriptional level and requires the products of the *Polycomb* and *piwi* genes in correlation with *Adh* promoter specific long non coding RNAs. These two proteins have been implicated in other silencing mechanisms that involve ncRNA molecules (Pal-Bhadra et al., 2002a, Malone et al., 2009). Because the silencing operates in trans, it is likely that a diffusible molecule is responsible for the recognition of all homologous copies. It is speculated that promoter specific long non coding RNA might sense all hybrid copies in the genome, which is required for optimal accumulation for transcriptional repressor proteins for unpaired DNA silencing.

In fact somatic pairing occurs in *Drosophila* cells allows the recognition of a similar but mechanistically distinct phenomenon. The silencing of unpaired DNA in multiple copies in the genome might have evolved as a sophisticated defense mechanism. A previous example of a copia LTR-CAT transgene exhibited silencing in the hemizygous state but not when homozygous (Matyunina et al., 2008). New transpositions of transposable elements and viral parasites will be present in an unpaired condition and thus might trigger a mechanism to silence the entire family of unpaired transpositions. The mode of action for unpaired DNA silencing is a novel example for transcriptional silencing in Drosophila. In Drosophila live cells the *Adh*-promoter w-reporter hybrid construct, mimics the behavior of different transposons and viral intruders. Thus initial insertions at the multiple sites of same *Adh-w* transgene transposition instigate the silencing event. Initially, promoter specific long non coding from the inserted element were generated. The long non coding RNA search the same transgene insertion copies in the genome. The Lnc RNA are coupled with the transgene promoter of the inserted copies which is required for proper docking for Pc and Piwi proteins. It eventually accumulates the histone repressor and its precursor protein EZ for proper and dramatic silencing in multiple copies. The silencing is also mediated by the histone modifications on the promoter, in particular, the methylation of H3me3Lys27. However, this relationship is likely complex because loss of the EZ methyltransferase does not completely abolish the H3 modification, suggesting that other methyltransferases (Cao and Zhang, 2004) are also recruited. The accumulation of Piwi is silenced transgene at a greatest magnitude and the release of EZ in the *piwi* mutants suggests an involvement in recruiting the EZ methyl-transferase for strong repression of the transgenes. Our findings also indicate that when an unpaired copy of *Adh-w* is present in the genome, Piwi becomes associated with the regulatory regions of the transgene (Figure 7). This association is coincident with the presence of the EZ histone methyltransferase and an increased H3me3K27 histone modification. This histone modification would be responsible for the strong transcriptional silencing. Moreover Lnc RNAs associated with piwi might serve as a homology guide to silence all copies in the genome including those that are paired, which alone do not exhibit such strong silencing. The Piwi in somatic ovarian follicle cells is implicated in transcriptional repression as the dominant mode of transposable element control in the soma (Ross et al., 2014). The transcriptional mechanism found here is coincident with chromatin modifications on the silenced transgenes.

The concept how unpaired DNA is recognized and repressed is not clear in many species. Many studies suggest that different mechanism may opt in different species (De Storme and Mason, 2014). At present Drosophila use a chromatin based mechanism alike Mammals that includes H3me3 K27 and other precursor proteins. In contrast N crassa adopted a post transcriptional mechanism (Aramayo and Selker, 2013). In *Neurospora crassa* DNA methylation is found at transposon and small repeats and depend on H3K9me2 but neither is dependent on RNAi and Piwi RNA biogenesis (Freitag et al., 2004). Nonetheless, two RNA silencing phenomena, quelling and meiotic silencing of unpaired DNA (MSUD) have been identified (Nakayashiki, 2005). MSUD senses and silences any foreign DNA that is unpaired during the homologue pairing stage of meiosis and all homologous copies of the gene, irrespectively (Lee et al., 2004).

A distinct set of silencing protein components seem to be involved in the MSUD and quelling pathways (Nakayashiki, 2005). However, neither DNA/histone methylation nor chromatin remodeling are implicated in MUSD (Shiu and Metzenberg, 2002) whereas at present, chromatin changes are implicated in neurospora. In *C. elegans*, EGO 1 and HIM-17 are scaffold proteins that may regulate heterochromatin. The H3me2K9 accumulation is disrupted in EGO-1 mutation. EGO1 also converts ssRNA tp dsRNA. The dsRNA without Dicer dependent cleavage relect nonprocessive RdRP activity. Similar to that chromatin modifying machinery also orchestrated different repressive molecules. In which Piwi functions as a non canonical manner. There may be certain protein yet to be identified that do not convert long non coding RNA to piRNA cluster. There may be ample precedent for involvement of long non coding RNA with Piwi proteins in the transcriptional repression (Rinn and Chang, 2012). It is not properly clear that which hybrid Adh-w gene behave differently, there is a reasonable limitation of white proteins in the adult fly. However presence of strong Adh promoter driven by white coding sequences produce ample of abnormal of white mRNA to neutralize the aberrant production of white proteins. Long non coding RNA was produced from the Adh promoter to reduce the transcriptional gene expression.

Indeed, Piwi and related Argonaute proteins are implicated in the control of transposons and viruses, most thoroughly studied in the germline(Brennecke et al., 2007, Siomi et al., 2008). Related functions of Piwi involve establishment of heterochromatin and telomere silencing (Pal-Bhadra et al., 2004b, Brower-Toland et al., 2007, Haynes et al., 2007, Yin and Lin, 2007, Josse et al., 2008). The function of piwi have also implicated the gene product in somatic silencing functions (Yin and Lin, 2007, Brower-Toland et al., 2007) including this study. The effect in somatic cells might arise from an early establishment of silencing during embryogenesis with perpetuation via the Polycomb complex or from a low level of expression in somatic cells.

A potential rationale for the existence of this strong silencing mechanism might be that it acts to counteract the expression of newly mobilized transposable elements (Matyunina et al., 2008). A new element insertion would by its nature be unpaired. A mechanism that recognizes the unpaired state of DNA in a novel context and strongly silences all homologous copies would serve as a genome defense against mutations created by new unpaired transposition. The involvement of a pairing component would be difficult to detect by analyzing multi-copy transposons; however, the use of defined transgenes allows recognition of this aspect of silencing (present study (Matyunina et al., 2008)). As noted above, silencing by unpaired DNA has been documented in meiosis in several taxa. Because Drosophila has somatic homologue pairing, this process can also occur in the soma as a protection against transposon activity and transposition.

## Experimental procedures

### Drosophila lines and genetic crosses

Flies were reared on *Drosophila* culture media at 25°C. The strains and mutations used in the experiments, if not noted, were described earlier (Pal-Bhadra et al., 1997, Pal-Bhadra et al., 1999) or in FLYBASE (http.flybase.org). The *Adh-w* hybrid transgenic lines were mobilized to different genomic locations by the presence of *P* element transposase provided by the *Δ2–3* strain (Robertson et al., 1988). Three different lines carrying the *Adh-w* construct located on separate chromosomes were used to generate the dosage series. The *Adh-w* insert on the X chromosome (*Adh-w#1*) is present at the 16B site. The *Adh-w#2* insert on the second chromosome was located to 55C2 and *Adh-w#3* on chromosome 3 is present at 70C.

To generate a series of (1-6 copies) *Adh-w* transgene stocks, we used multiple balancer chromosomes to combine different doses of the *Adh-w* transgene. The design of each cross is described in Supplementary Materials.

### Eye pigment assay

For pigment quantification, forty eight adult heads for each genotype were dissected manually, collected, homogenized in 1.0 ml of methanol acidified with 0.1% HCl and subjected to centrifugation. The supernatant was used to measure absorbance at 480 nm.

### RNA isolation and Northern blot hybridization

To estimate the *white* mRNA level from each genotype, total cellular RNA extracted from adult flies was denatured, subjected to electrophoresis in agarose gels, transferred to nylon membrane and hybridized with antisense radio-labeled *white* RNA and counter hybridized with antisense *β-tubulin* probes as described earlier (Pal-Bhadra et al., 1997). The amount of radioactivity in each band was estimated using a Fuji phosphorimager.

### RNA protection assay

Extraction of cellular RNA, *in vitro* RNA transcription, hybridization, RNase digestion and acrylamide gel electrophoresis were performed as per the manufacturer's instructions with minor modifications. RNA hybrids were digested at 37°C for 25 minutes with 1:150 dilution of RNase T1 stock solution for the probe. The product was fractionated on 10% denaturing polyacrylamide gels.

### Nuclear run on transcription

Adult flies were collected and used for isolating fresh nuclei for transcriptional run-on experiments as described (So and Rosbash, 1997). The estimate of the amount of *white* DNA that is required for complete hybridization with the RNA generated from 6 copies of the *Adh-w* transgene, a dilution series of *Adh-w* DNA was used to determine the maximal amount of DNA under these circumstances. The experiment determined that nearly 10 ug of *white* DNA was required to prevent saturation of RNA hybridization (**Figure S8**).

### Chromosomal immunostaining

Preparation, fixation and immunostaining of the polytene chromosomes from the salivary glands of third instar larvae were performed as described (Pal-Bhadra et al., 2004b). The chromosomes were probed with rat anti-Polycomb (1:200) antibodies. For confocal microscopy Cy-5 conjugated goat anti-rabbit (1:200), and FITC conjugated goat anti-rat (1:200) antibodies were used. After incubation, the chromosomes were thoroughly washed with buffer (400 mM NaCl, 0.2% Tween 20, 0.2% NP40 in PBS) to remove background. The slides were mounted with DAPI or propidium iodide containing antifade Vectashield mounting media and examined with an Olympus Fv1000 confocal microscope using a 60X oil lens.

The chromosomes were also immunostained with commercially available rabbit anti-PIWI (Abcam batch # ab 5207-100) antibodies with a standard concentration (1:100) at which there was no detectable binding. Subsequently, anti-PIWI antibodies were used at a higher concentration (1:50, 1: 25, and 1:10) (**Figure S4**). Association of anti-PIWI antibodies with chromosomes was proportionately increased with the concentration and maximal at the 1:10 dilution (**See** Figure 5). For double immunostaining, we used the same concentration of rabbit polyclonal anti-PIWI antibody (1:10) to show its overlap with anti-PC antibody. To verify that the antibody is specific to PIWI protein, the chromosomes from heteroallelic *piwi (piwf/piwi^2^)* mutant larvae were immunostained at the same concentration (1:10 dilution) (**Figure S4**).

Several previous reports showed that PIWI is associated with a novel species of RNA (piRNA) and other non-coding RNAs (Brennecke et al., 2007, Gunawardane et al., 2007). It is reasonable to believe that RNAs might associate with PIWI and create binding sites for anti-PIWI on the chromosomes. To prevent the degradation of RNA, we omitted an RNase treatment of the chromosomes when PIWI antibodies were used. As a result, the clarity of the chromosomal binding is reduced, which is compromised in double and anti-PIWI immunostained chromosome preparations.

### RNA immunoprecipitation (RIP)

The sheared 200 μg of cross-linked chromatin was diluted with ChIP dilution buffer [0.01% SDS, 1.1% Triton-X100, 1.2 mM EDTA, 16.7 mM Tris-HCl pH 8.1, 167 mM NaCl, 1X EDTA free protease inhibitors cocktail (Roche), RNase out] and pre-cleared with 50% Protein A agarose/salmon sperm DNA beads (Millipore) for 1 hour at 4°C. Then 150 - 200 μg of pre-cleared chromatin was incubated with 2-5 μg of antibodies overnight prior to the addition of a protein A Sepharose beads. Immunoprecipitated complexes were washed sequentially with low salt buffer (0.1% SDS, 1% Triton X100, 2 mM EDTA, 20 mM Tris-HCl pH8.1, 150 mM NaCl), high salt buffer (0.1% SDS, 1% Triton-X100, 2 mM EDTA, 20 mM Tris-HCl pH8.1, 500 mM NaCl), LiCl wash buffer (0.25 M LiCl, 1% NP-40, 1% deoxycholate, 1 mM EDTA, 10 mM Tris-HCl pH8.1) and 10 mM Tris-HCl, 1 mM EDTA pH8 (2 times). The bound DNA was eluted with elution buffer (1% SDS, 0.1 M Na_2_HCO3) and cross-links removed at 65°C for 2 h. After treatment with Proteinase K (Sigma) for 45 min at 42°C, RIP RNA was isolated following standard Trizol method (Invitrogen). Then RNA was treated with TURBO DNase 1 (Ambion) to remove DNA contamination. Single stranded cDNA was synthesized using random hexamer by Superscript II RT kit (Invitrogen). Then the amplification was performed with following *mini-w* promoter specific primer sets (previously mentioned) at 55°C annealing temperature.

### Chromatin Immuno-precipitation (ChIP)

To detect the amount of POLYCOMB, EZ and histone H3K27me3 proteins associated with the regulatory region of the *Adh-w* transgene, chromatin immuno-precipitation was conducted as described previously (Cavalli and Paro, 1999). Cells were cross-linked with formaldehyde, fixed chromatin washed extensively, reverse cross-linked, and digested with RNase A and proteinase K. The immuno-precipitated DNA was amplified by semi-quantitative PCR and real time PCR using primers specific for the *Adh* promoter and *white* exon 2. The coding region of the *white* genes was amplified as an internal control (Figure 7). We also performed ChiP analysis using PIWI antibodies (Abcam batch # ab 5207-100). For estimation of antibodies, immuno-precipitated and input DNA were amplified in the same reaction for 35 cycles at 94^o^C for 45 sec, 58^o^C for 45 sec, 72^o^C for 1 min using two primer pairs spanning different promoter regions. The PCR products were electrophoresed in a 2% agarose gel. The following oligonucleotides were used: *Adh* promoter forward primer TCCAACTTTTCTAGATTGATTC, reverse primer ATCTGCAAACAAATGCGGGGT; *white* second exon forward primer TCGCAGAGCTGCATTAACC, reverse primer ATTGACCGCCCCAAAGAT.

## Acknowledgments

We thank V. Pirrotta and the *Drosophila* Stock Center at Indiana University, USA for supplying fly stocks, G. Cavalli, R. Paro and Greg Hannon for antibodies and I. Ganguly and I. Bag for helping with the inverse PCR method and polytene staining. This work was supported by a Wellcome Trust Senior Research Fellowship (GRA070065MA to UB and GRA076395AIA to MPB), HFSP Young investigator grant (RGY20/2003).

## References

Aramayo, R. & Selker, E. U. 2013. Neurospora crassa, a model system for epigenetics research. Cold Spring Harb Perspect Biol, 5, a017921.

Aravin, A. & Tuschl, T. 2005. Identification and characterization of small RNAs involved in RNA silencing. FEBS Lett, 579, 5830–40.

Aravin, A. A., Hannon, G. J. & Brennecke, J. 2007. The Piwi-piRNA pathway provides an adaptive defense in the transposon arms race. Science, 318, 761–4.

Baarends, W. M., Wassenaar, E., Van Der Laan, R., Hoogerbrugge, J., Sleddens-Linkels, E., Hoeijmakers, J. H., De Boer, P. & Grootegoed, J. A. 2005. Silencing of unpaired chromatin and histone H2A ubiquitination in mammalian meiosis. Mol Cell Biol, 25, 1041–53.

Berry, B., Deddouche, S., Kirschner, D., Imler, J. L. & Antoniewski, C. 2009. Viral suppressors of RNA silencing hinder exogenous and endogenous small RNA pathways in Drosophila. PLoS One, 4, e5866.

Brennecke, J., Aravin, A. A., Stark, A., Dus, M., Kellis, M., Sachidanandam, R. & Hannon, G. J. 2007. Discrete small RNA-generating loci as master regulators of transposon activity in Drosophila. Cell, 128,1089–103.

Brower-Toland, B., Findley, S. D., Jiang, L., Liu, L., Yin, H., Dus, M., Zhou, P., Elgin, S. C. & Lin, H. 2007. Drosophila PIWI associates with chromatin and interacts directly with HP1a. Genes Dev, 21, 2300–11.

Cao, R. & Zhang, Y. 2004. SUZ12 is required for both the histone methyltransferase activity and the silencing function of the EED-EZH2 complex. Mol Cell, 15, 57–67.

Cavalli, G. & Paro, R. 1999. Epigenetic inheritance of active chromatin after removal of the main transactivator. Science, 286, 955–8.

Chung, W. J., Okamura, K., Martin, R. & Lai, E. C. 2008. Endogenous RNA interference provides a somatic defense against Drosophila transposons. Curr Biol, 18, 795–802.

Corbin, V. & Maniatis, T. 1989a. The role of specific enhancer-promoter interactions in the Drosophila Adh promoter switch. Genes Dev, 3, 2191–20.

Corbin, V. & Maniatis, T. 1989b. Role of transcriptional interference in the Drosophila melanogaster Adh promoter switch. Nature, 337, 279–82.

Cox, D. N., Chao, A. & Lin, H. 2000. piwi encodes a nucleoplasmic factor whose activity modulates the number and division rate of germline stem cells. Development, 127, 503–14.

De Storme, N. & Mason, A. 2014. Plant speciation through chromosome instability and ploidy change: Cellular mechanisms, molecular factors and evolutionary relevance. Current Plant Biology, 1, 10–33.

Fagegaltier, D., Bouge, A. L., Berry, B., Poisot, E., Sismeiro, O., Coppee, J. Y., Theodore, L., Voinnet, O. & Antoniewski, C. 2009. The endogenous siRNA pathway is involved in heterochromatin formation in Drosophila. Proc Natl Acad Sci U S A, 106, 21258–63.

Franke, A., Decamillis, M., Zink, D., Cheng, N., Brock, H. W. & Paro, R. 1992. Polycomb and polyhomeotic are constituents of a multimeric protein complex in chromatin of Drosophila melanogaster. EMBO J, 11, 2941–50.

Freitag, M., Lee, D. W., Kothe, G. O., Pratt, R. J., Aramayo, R. & Selker, E. U. 2004. DNA methylation is independent of RNA interference in Neurospora. Science, 304, 1939.

Ghildiyal, M., Seitz, H., Horwich, M. D., Li, C., Du, T., Lee, S., Xu, J., Kittler, E. L., Zapp, M. L., Weng, Z. & Zamore, P. D. 2008. Endogenous siRNAs derived from transposons and mRNAs in Drosophila somatic cells. Science, 320, 1077–81.

Grewal, S. I. & Elgin, S. C. 2007. Transcription and RNA interference in the formation of heterochromatin. Nature, 447, 399–406.

Grimaud, C., Bantignies, F., Pal-Bhadra, M., Ghana, P., Bhadra, U. & Cavalli, G. 2006. RNAi components are required for nuclear clustering of Polycomb group response elements. Cell, 124, 957–71.

Gunawardane, L. S., Saito, K., Nishida, K. M., Miyoshi, K., Kawamura, Y., Nagami, T., Siomi, H. & Siomi, M. C. 2007. A slicer-mediated mechanism for repeat-associated siRNA 5′ end formation in Drosophila. Science, 315,1587–90.

Haynes, K. A., Gracheva, E. & Elgin, S. C. 2007. A Distinct type of heterochromatin within Drosophila melanogaster chromosome 4. Genetics, 175, 1539–42.

Hazelrigg, T., Levis, R. & Rubin, G. M. 1984. Transformation of white locus DNA in drosophila: dosage compensation, zeste interaction, and position effects. Cell, 36, 469–81.

Hynes, M. J. & Todd, R. B. 2003. Detection of unpaired DNA at meiosis results in RNA-mediated silencing. Bioessays, 25, 99–103.

Josse, T., Maurel-Zaffran, C., De Vanssay, A., Teysset, L., Todeschini, A. L., Delmarre, V., Chaminade, N., Anxolabehere, D. & Ronsseray, S. 2008. Telomeric trans-silencing in Drosophila melanogaster: tissue specificity, development and functional interactions between non-homologous telomeres. PLoS One, 3, e3249.

Lee, D. W., Seong, K.-Y., Pratt, R. J., Baker, K. & Aramayo, R. 2004. Properties of unpaired DNA required for efficient silencing in Neurospora crassa. Genetics, 167, 131–150.

Malone, C. D., Brennecke, J., Dus, M., Stark, A., Mccombie, W. R., Sachidanandam, R. & Hannon, G. J. 2009. Specialized piRNA pathways act in germline and somatic tissues of the Drosophila ovary. Cell, 137, 522–35.

Matyunina, L. V., Bowen, N. J. & Mcdonald, J. F. 2008. LTR retrotransposons and the evolution of dosage compensation in Drosophila. BMC Mol Biol, 9, 55.

Moazed, D. 2009. Small RNAs in transcriptional gene silencing and genome defence. Nature, 457, 413–20.

Nakayashiki, H. 2005. RNA silencing in fungi: mechanisms and applications. FEBS Lett, 579, 5950–7.

Pal-Bhadra, M., Bhadra, U. & Birchler, J. A. 1997. Cosuppression in Drosophila: gene silencing of Alcohol dehydrogenase by white-Adh transgenes is Polycomb dependent. Cell, 90, 479–90.

Pal-Bhadra, M., Bhadra, U. & Birchler, J. A. 1999. Cosuppression of nonhomologous transgenes in Drosophila involves mutually related endogenous sequences. Cell, 99, 35–46.

Pal-Bhadra, M., Bhadra, U. & Birchler, J. A. 2002a. RNAi related mechanisms affect both transcriptional and posttranscriptional transgene silencing in Drosophila. Mol Cell, 9, 315–27.

Pal-Bhadra, M., Bhadra, U. & Birchler, J. A. 2002b. RNAi related mechanisms affect both transcriptional and posttranscriptional transgene silencing in Drosophila. Molecular cell, 9, 315–327.

Pal-Bhadra, M., Bhadra, U. & Birchler, J. A. 2004a. Interrelationship of RNA interference and transcriptional gene silencing in Drosophila. Cold Spring Harb Symp Quant Biol, 69, 433–8.

Pal-Bhadra, M., Leibovitch, B. A., Gandhi, S. G., Chikka, M. R., Bhadra, U., Birchler, J. A. & Elgin, S. C. 2004b. Heterochromatic silencing and HP1 localization in Drosophila are dependent on the RNAi machinery. Science, 303, 669–72.

Palm, W., Sampaio, J. L., Brankatschk, M., Carvalho, M., Mahmoud, A., Shevchenko, A. & Eaton, S. 2012. Lipoproteins in Drosophila melanogaster––assembly, function, and influence on tissue lipid composition. PLoS Genet, 8, e1002828.

Rastelli, L., Chan, C. S. & Pirrotta, V. 1993. Related chromosome binding sites for zeste, suppressors of zeste and Polycomb group proteins in Drosophila and their dependence on Enhancer of zeste function. EMBO J, 12, 1513–22.

Rinn, J. L. & Chang, H. Y. 2012. Genome regulation by long noncoding RNAs. Annu Rev Biochem, 81, 145–66.

Robertson, H. M., Preston, C. R., Phillis, R. W., Johnson-Schlitz, D. M., Benz, W. K. & Engels, W. R. 1988. A stable genomic source of P element transposase in Drosophila melanogaster. Genetics, 118, 461–70.

Ross, R. J., Weiner, M. M. & Lin, H. 2014. PIWI proteins and PIWI-interacting RNAs in the soma. Nature, 505, 353–9.

She, X., Xu, X., Fedotov, A., Kelly, W. G. & Maine, E. M. 2009. Regulation of heterochromatin assembly on unpaired chromosomes during Caenorhabditis elegans meiosis by components of a small RNA-mediated pathway. PLoS Genet:, 5, e1000624.

Shiu, P. K. & Metzenberg, R. L. 2002. Meiotic silencing by unpaired DNA: properties, regulation and suppression. Genetics, 161, 1483–95.

Shiu, P. K., Raju, N. B., Zickler, D. & Metzenberg, R. L. 2001. Meiotic silencing by unpaired DNA. Cell, 107, 905–16.

Siomi, M. C., Saito, K. & Siomi, H. 2008. How selfish retrotransposons are silenced in Drosophila germline and somatic cells. FEBS Lett, 582, 2473–8.

So, W. V. & Rosbash, M. 1997. Post-transcriptional regulation contributes to Drosophila clock gene mRNA cycling. EMBO J, 16, 7146–55.

Steller, H. & Pirrotta, V. 1985. Expression of the Drosophila white gene under the control of the hsp70 heat shock promoter. EMBO J, 4, 3765–72.

Sullivan, W., Ashburner, M. & Hawley, R. S. 2000. Drosophila Protocols, Cold Spring Harbor Laboratory Press.

Thomson, T. & Lin, H. 2009. The biogenesis and function of PIWI proteins and piRNAs: progress and prospect. Annu Rev Cell Dev Biol, 25, 355–76.

Volpe, T. A., Kidner, C., Hall, I. M., Teng, G., Grewal, S. I. & Martienssen, R. A. 2002. Regulation of heterochromatic silencing and histone H3 lysine-9 methylation by RNAi. Science, 297, 1833–7.

Yin, H. & Lin, H. 2007. An epigenetic activation role of Piwi and a Piwi-associated piRNA in Drosophila melanogaster. Nature, 450, 304–8.

